# Combining Backpropagation with Equilibrium Propagation to improve an Actor-Critic RL framework

**DOI:** 10.1101/2022.06.21.496871

**Authors:** Yoshimasa Kubo, Eric Chalmers, Artur Luczak

## Abstract

Backpropagation has been used to train neural networks for many years, allowing them to solve a wide variety of tasks like image classification, speech recognition, and reinforcement learning tasks. But the biological plausibility of backpropagation as a mechanism of neural learning has been questioned. Equilibrium Propagation (EP) has been proposed as a more biologically plausible alternative and achieves comparable accuracy on the CIFAR-10 image classification task. This study proposes the first EP-based reinforcement learning architecture: an actor-critic architecture with the actor network trained by EP. We show that this model can solve the basic control tasks often used as benchmarks for BP-based models. Interestingly, our trained model demonstrates more consistent high-reward behavior than a comparable model trained exclusively by backpropagation.

## Introduction

The backpropagation (BP) algorithm [1] has long been the workhorse of deep neural networks, allowing their successful application to many tasks. BP-powered neural networks have enabled reinforcement learning systems to outperform humans at Go [2] and Atari games [3]. But BP has been criticized as being not biologically plausible (it seems unlikely that neurons do anything like compute partial derivatives). It has also been observed that humans still outperform deep neural networks on many tasks, like adversarial examples [4], art and music. Could more biologically-plausible learning mechanisms help close this gap?

In the reinforcement learning context, one biologically-plausible method is the REINFORCE framework - a policy-gradient algorithm that was described in a neuroscience context by Williams [5]. The parallels between REINFORCE and biological neural learning have been discussed by Sutton et al [6] and Chung [7], and it has led to more recent developments such as the Attention-Gated Brain Propagation approach [8]. Actor-Critic is another reinforcement learning architecture with parallels to biological learning: several studies have seen the actor-critic architecture as an analog of learning mechanisms in the basal ganglia [9-11]. Biologically-plausible reinforcement learning approaches can demonstrate more human-like behavior [12], and so may provide important insights into human learning and intelligence.

In the supervised learning context, Equilibrium Propagation (EP) has been proposed as a more biologically plausible alternative to backpropagation [13-17]. EP is an extension of Contrastive Hebbian Learning [18-20] that sees the neural network as a dynamical system whose steady state can be perturbed by inputs during an initial “free” phase, and then clamped by teaching signals in a second “clamped” phase, affecting learning in a biologically realistic way. EP has successfully trained algorithms to perform image classification tasks like MNIST [21] and CIFAR10 [22], and Laborieux et al showed that convolutional networks trained by EP can achieve comparable accuracy to backpropagation in the CIFAR10 task [14]. A further extension of EP by Luczak et al. showed how learning might occur in a single phase - making the algorithm even more biologically plausible - while still achieving good classification accuracy [23].

A biologically plausible reinforcement learning approach based on EP has not yet been proposed. Here we explore an Actor-Critic architecture trained by both backpropagation and by brain-inspired modification of EP proposed by Luczak et al [23]. This study provides two contributions:

1. We propose the first application of Equilibrium Propagation to reinforcement learning, in the form of an Actor-Critic architecture trained by a combination of EP (Actor) and backpropagation (Critic).
2. We demonstrate that our architecture can solve several control tasks, and that its learned behaviors are more consistently rewarding than behaviors learned using backpropagation alone.

## Method

This section details how our Actor-Critic architecture was implemented.

### Actor Critic Architecture

Actor-Critic is a two-part architecture for reinforcement learning. The “Actor” is a model that encapsulates the learner’s policy: it observes the current state and outputs an action to execute. The “Critic” is a separate model that estimates the value of an action given a particular state. It observes the effect of each executed action, often in the form of a difference between the predicted value of the action and the value actually experienced (a “temporal difference error”). It uses the temporal difference error as a learning signal to improve its own future value estimates, and also to update the Actor to make high-value actions more likely, and low-value actions less likely.

### Actor network (trained by Equilibrium Propagation)

EP envisions a neural network as a dynamical system that learns in two phases. First is the “free phase”, in which an input is applied and the network is allowed to equilibrate. During this phase the network dynamics obey the equations:

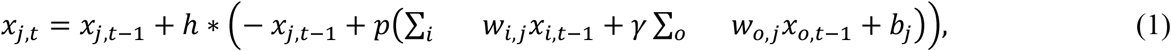

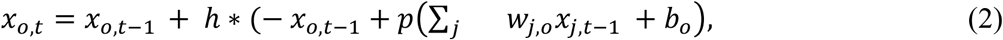

where *x* is an activation, *w* is weights for each layer, *i, j*, and *o*, are indexes of input, hidden and output layer neurons, *b* is a bias. *P* is an activation function such as the sigmoid function, and *γ* is the feedback parameter. *h* is the Euler method’s time-step.

After the network has reached a free-phase steady state, the second “clamped” phase begins. During this phase the output neurons are clamped (or rather, weakly clamped or nudged) toward the target values. In conventional EP the dynamics during this phase obey the equations:

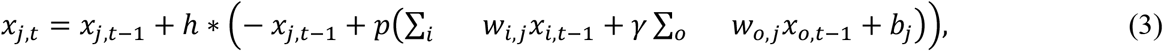

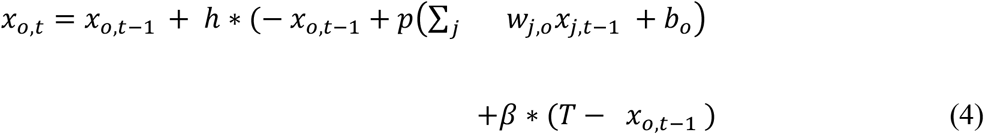

where *T* is a target for the classification task.

However, in a reinforcement learning setting there is no target signal per se; only the reward signal, which the learner must use to estimate values of particular states and actions in the environment. To accommodate this different paradigm, our Actor network modifies equation (4) as follows:

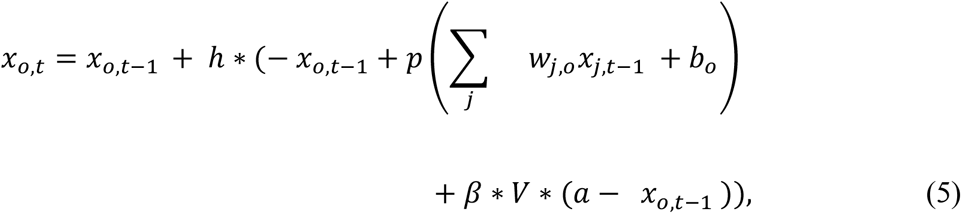

where *a* is the action that was taken, and *V* is the estimated value of the state, as estimated by the critic network. Alternatively, *V* can be replaced with a temporal-difference-style quantity:

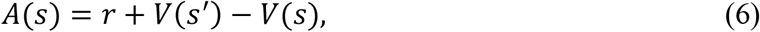

Where *s* is the current state, and *s’* is the new state (arrived at after executing *a*). This makes the clamped-phase dynamics at the output:

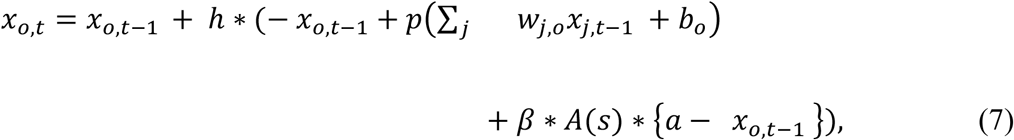

After the network reaches a clamped-phase steady state, weights could be updated according to the rule derived in the original EP paper [16]:

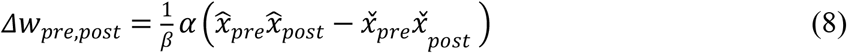

where 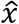 is an activity at the weakly clamped phase, 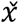 is an activity at the free phase, *α* is the learning rate, *β* is a nudging parameter, *pre* and *post* are previous and post layer neuron indexes, respectively (e.g. for *Δw*_*i,j*_, *pre* and *post* will be *i* and *j*, respectively).

Here we replace equation (8) with the new rule proposed in our previous work [23], which allows learning to occur in a single phase by assuming that neurons may predict their own future activity.

The study showed that a rule of this form emerges naturally if we assume that each neuron is working to maximize its metabolic energy. The new rule is:

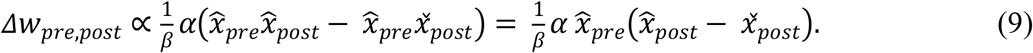

### Critic network (trained by backpropagation)

Equation 5 represents a prediction error - the error between *r* + *V*(*s’*), the actual value of the present experience, and *V*(*s*), the predicted value. The mean squared prediction error is then:

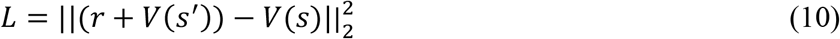

The critic network is tuned using backpropagation in the usual way to minimize this prediction error.

### Experience replay

We use experience replay [3, 24-26] to make our model more stable. This method stores the agent’s experiences (including states, actions, rewards, and next-states) and makes them available for learning later. It is worth noting that experience replay is also biologically plausible; analogous to memory replay during sleep [27].

The complete algorithm is shown in Algorithm 1.

#### Algorithm 1

Train Actor-Critic by EP and BP.

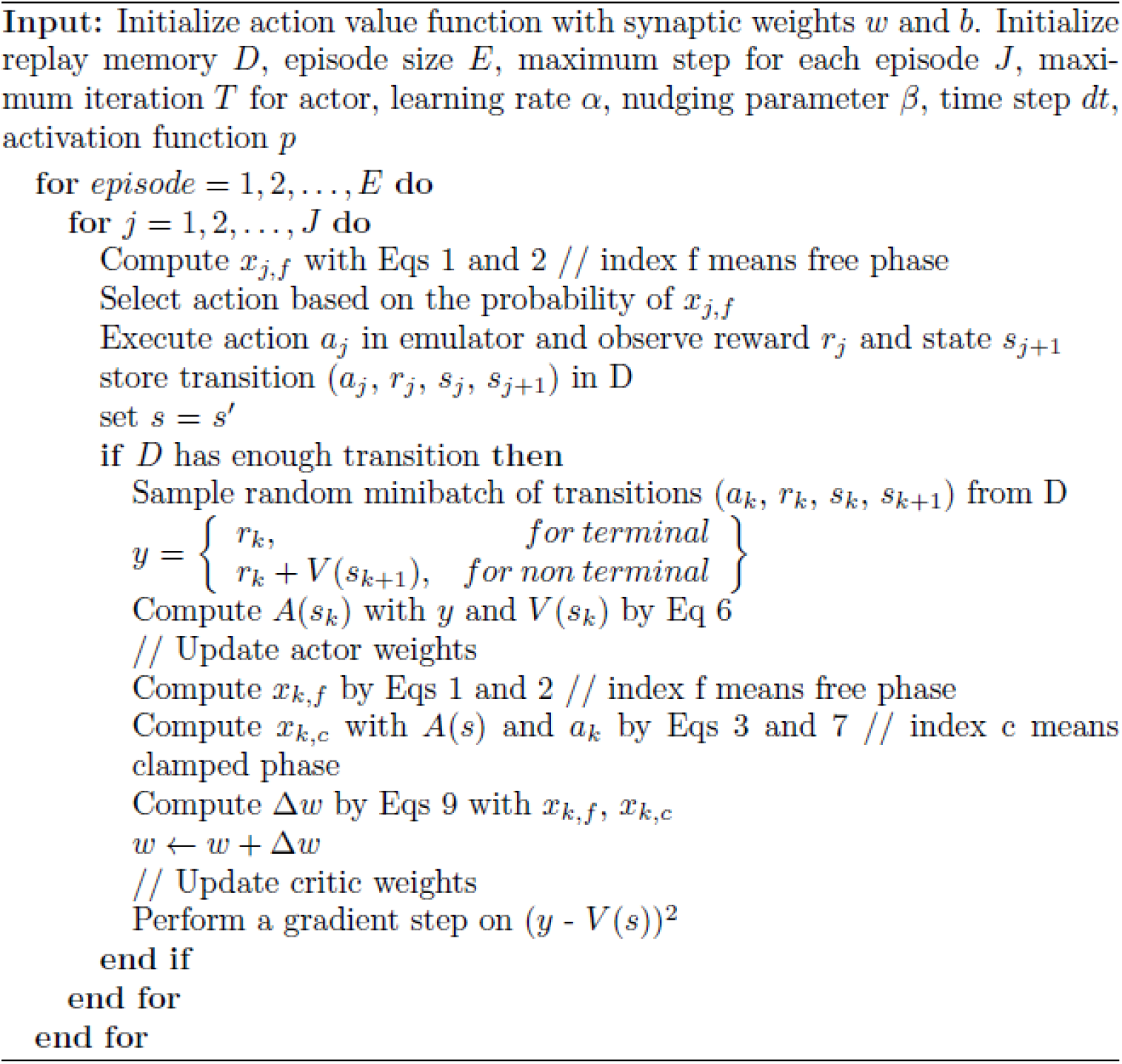

The code to reproduce our results is located at:

https://github.com/ykubo82/HybridRL

## Experiments

We tested our model in three simple Open AI gym tasks [28]: CartPole-v0, Acrobot-v1, and LunarLander-v2. All of these tasks feature continuous states and discrete actions. Our model uses multilayer perceptrons for both Actor and Critic networks, trained by EP and backpropagation, respectively. Each MLP consists of 1 hidden layer with 256 nodes. For LunarLander-v2, we increased the hidden size to 512 due to the complexity of the task.

The activation function for the hidden layer on both Actor and Critic is the hard sigmoid from [14] for CartPole-v0 and LunarLander-v2, the hard sigmoid from [13] for Acrobot-v1. The activation function for the Actor’s output layer is the softmax function. A maximum of 1000 experiences were stored for experience replay, and the mini-batch size was 20. The learner was allowed 1000 steps for the CartPole-v0 and Acrobot-v1 tasks, and 2000 for LunarLander-v2. Parameter settings are shown in Table 1. For critic networks trained by backpropagation, we used Adam optimizer [29] to accelerate models’ training.

**Table 1:**
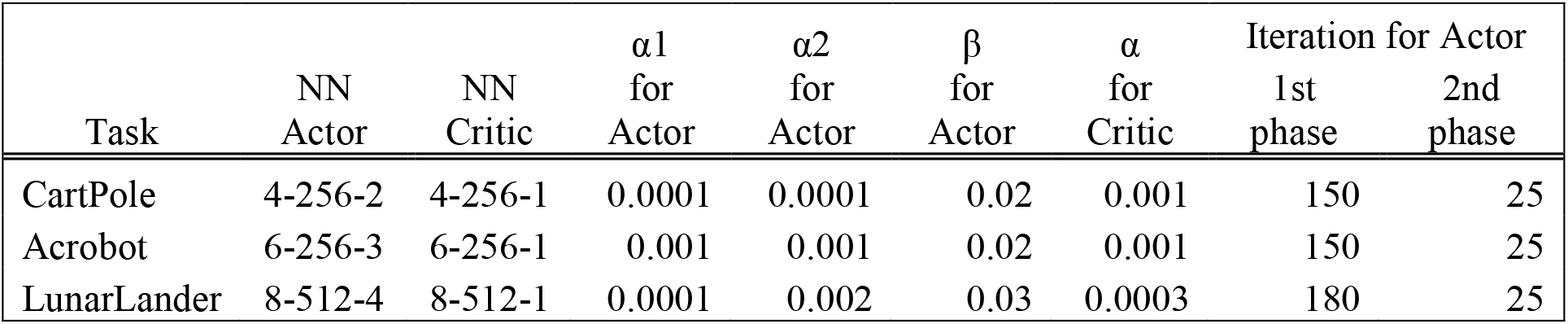
Parameters for our models on each task. NN describes number of neurons in each layer, α_1_ is the learning rate for the weights between the input and hidden layer, α_2_ is the learning rate for the weights between the hidden and output layers, and 1st and 2nd phases mean duration of free phase and weakly clamped phases, respectively.

For comparison, we also implement a model with the same architecture as described above, but trained purely by backpropagation. Hereafter we refer to our model with Actor trained by EP and Critic trained by backpropagation as EP-BP, while the baseline Actor-Critic model trained entirely by backpropagation as BP. All models were run 8 times, and means and standard deviations were recorded.

## Results

Figure 2 shows performance of EP-BP and BP on each task. On all tasks our EP-BP model converges to more stable rewarding behavior than the baseline model train with BP only. This is quantified in Figures 3 which shows the mean reward obtained in the last 25% episodes. In each case the mean reward obtained by EP-BP is higher as compared to BP model. Moreover, closer examination of traces in Figure 2 showed higher variability in reward for BP trained model. To quantify it, for each of 8 runs of the model we calculated SD from the last 25% of episodes. Figure 4 shows average SD across 8 runs for each model. This measure of variability was consistently lower for our EP-BP model. This tells us that our model is more stable than the base line model.

**Figure 1:**
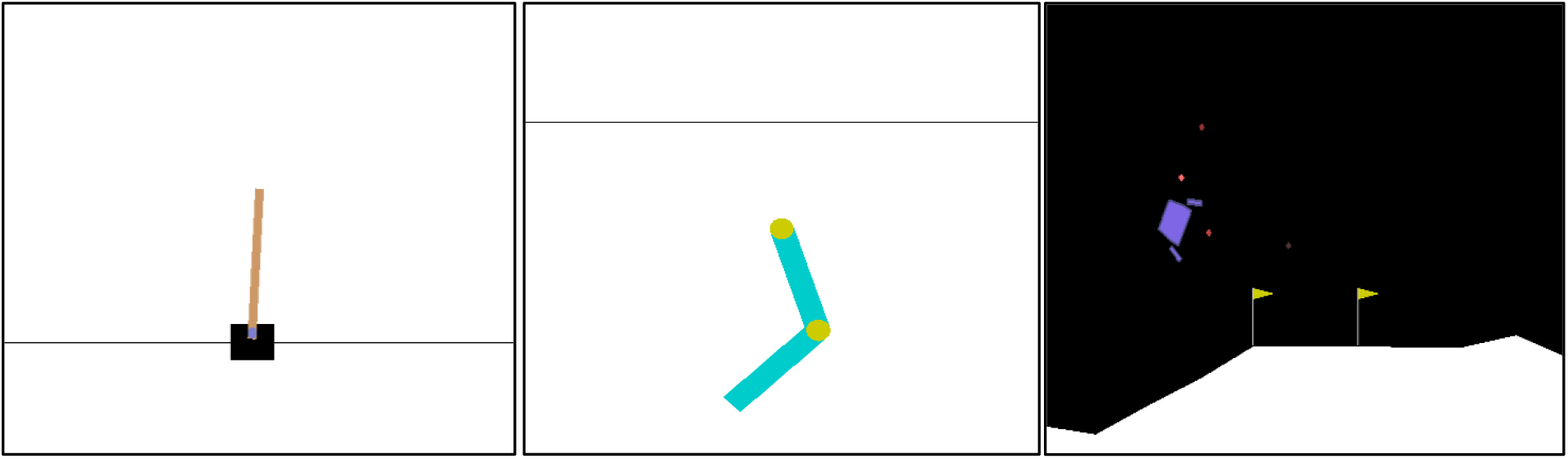
Images of environments in tasks for our model: CartPole-v0 (left), Acrobot-v1 (center), and LunarLander-v2 (right). CartPole-v0 task: A pole is on a cart, and this pole is unstable. The goal of this task is to move the cart to left or right to balance the pole. Acrobot-v1: a robot arm is composed of two joints. The goal of this task is to swing the arm to reach the black horizontal line. LunarLander-v2: There is a spaceship that tries to land. The goal of this task is to land the spaceship between the flags smoothly by moving the spaceship.

**Figure 2:**
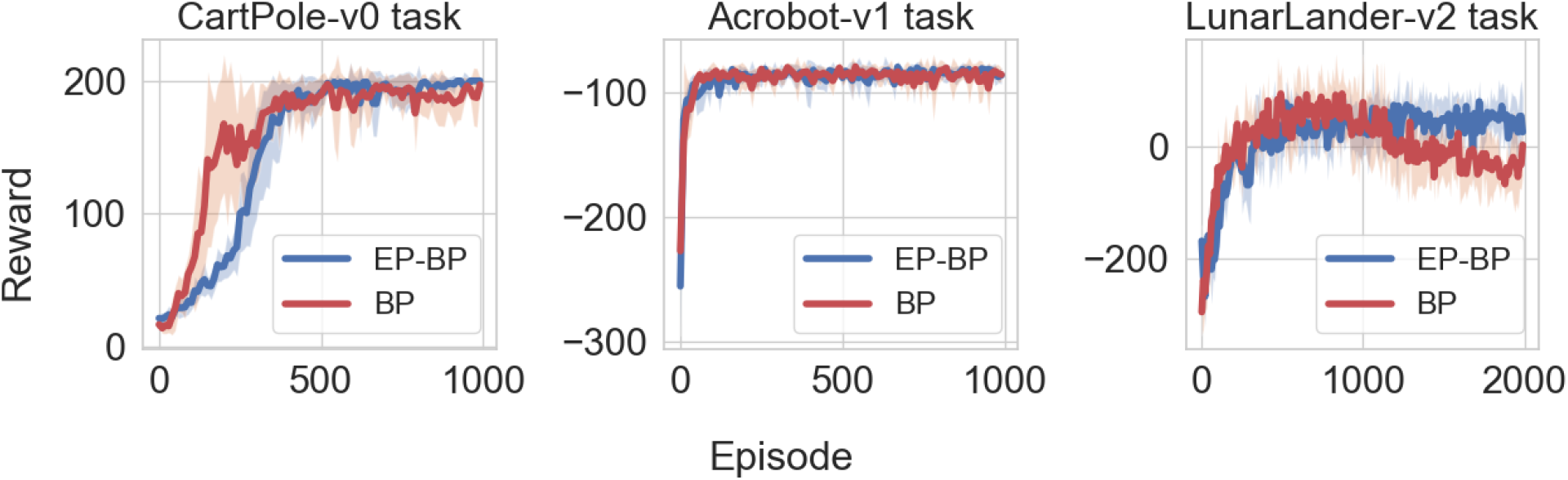
Plotting the reward vs episode for CartPole-v0 (top-left), Acrobot-v1 (top-right), and LunarLander-v2 (bottom) on both BP and EP-BP. Solid lines shows mean across 8 runs and shaded area denote standard deviation. Note that for Acrobot-v1, the agent receives -1 as punishment until it reaches the target.

**Figure 3:**
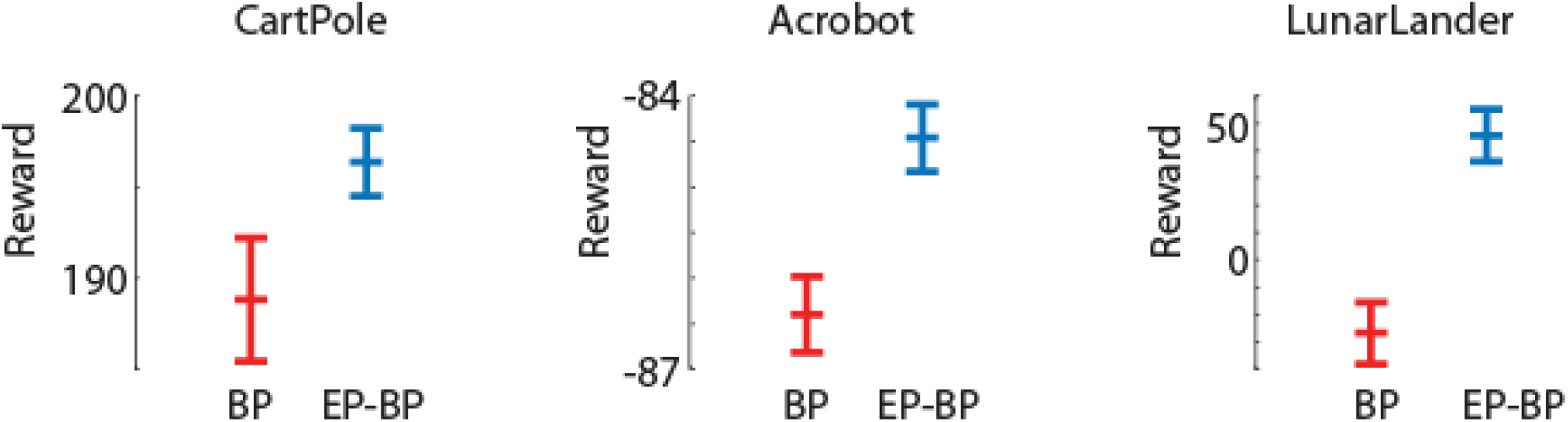
Average rewards and std error (SEM) for the last 25% of episodes for BP and EP-BP on CartPole-v0, Acrobot-v1, and LunarLander-v2.

**Figure 4:**
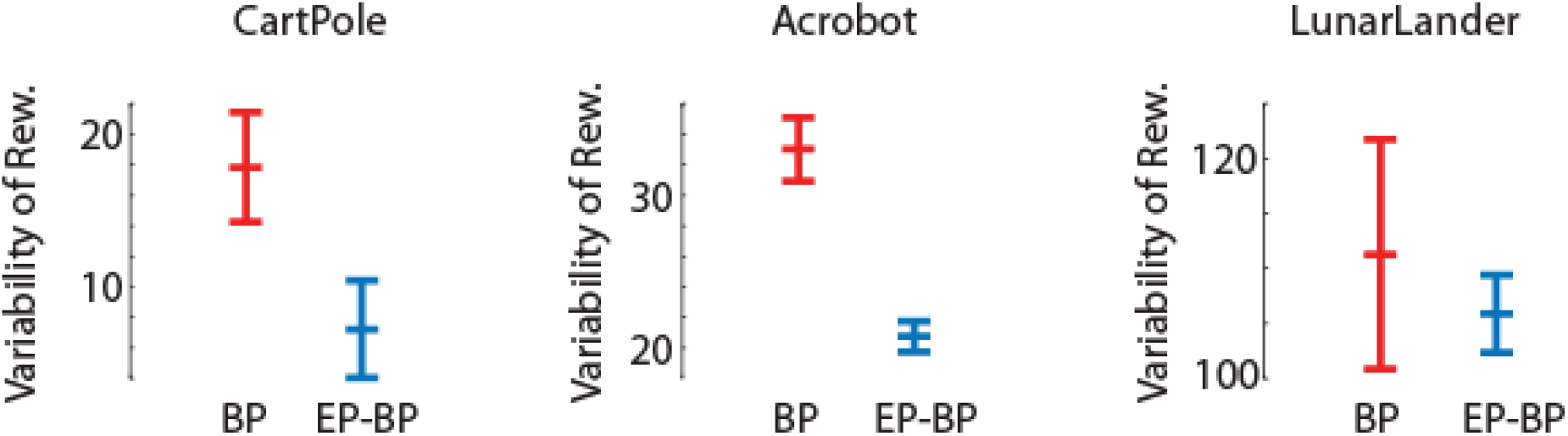
Average variability and std error (SEM) for the last 25% of episodes for BP and EP-BP on CartPole-v0, Acrobot-v1, and LunarLander-v2.

As an internal measure of learning, similar to Römer et al [30], we recorded the softmax probability for each action executed by the Actor network throughout learning. For each episode, we saved the probabilities of actions that EP-BP and BP took. Figure 5 shows EP-BP executing actions with very high (>90%) confidence after about 600 episodes in the CarPole-v0 task. This means that less than 10% of actions are selected randomly. Randomness might be important for exploring the environment in the early phase for gathering information about the environment (exploration), but in the last phase, the model should take the optimal action after getting enough information (exploitation) [31]. However, if a model does not have a high enough confidence which action is optimal, the model might not take that action because there is still some randomness. For example, a person knows that A route is always busy with traffic jams based on his experience (thanks to exploration), thus he always takes B route to the office and arrives on time (exploitation). However, another person also knows that A route is always busy based on his experience, but he sometimes takes the A route (more often than the first person) because he does not have enough confidence for the B route (this means he thinks sometimes the B route might not be busy), and he is sometimes late. Thus the BP model’s confidence is lower after learning, which may explain its somewhat less consistent behaviour (this means the BP model takes more often non-optimal actions than EP-BP model).

**Figure 5:**
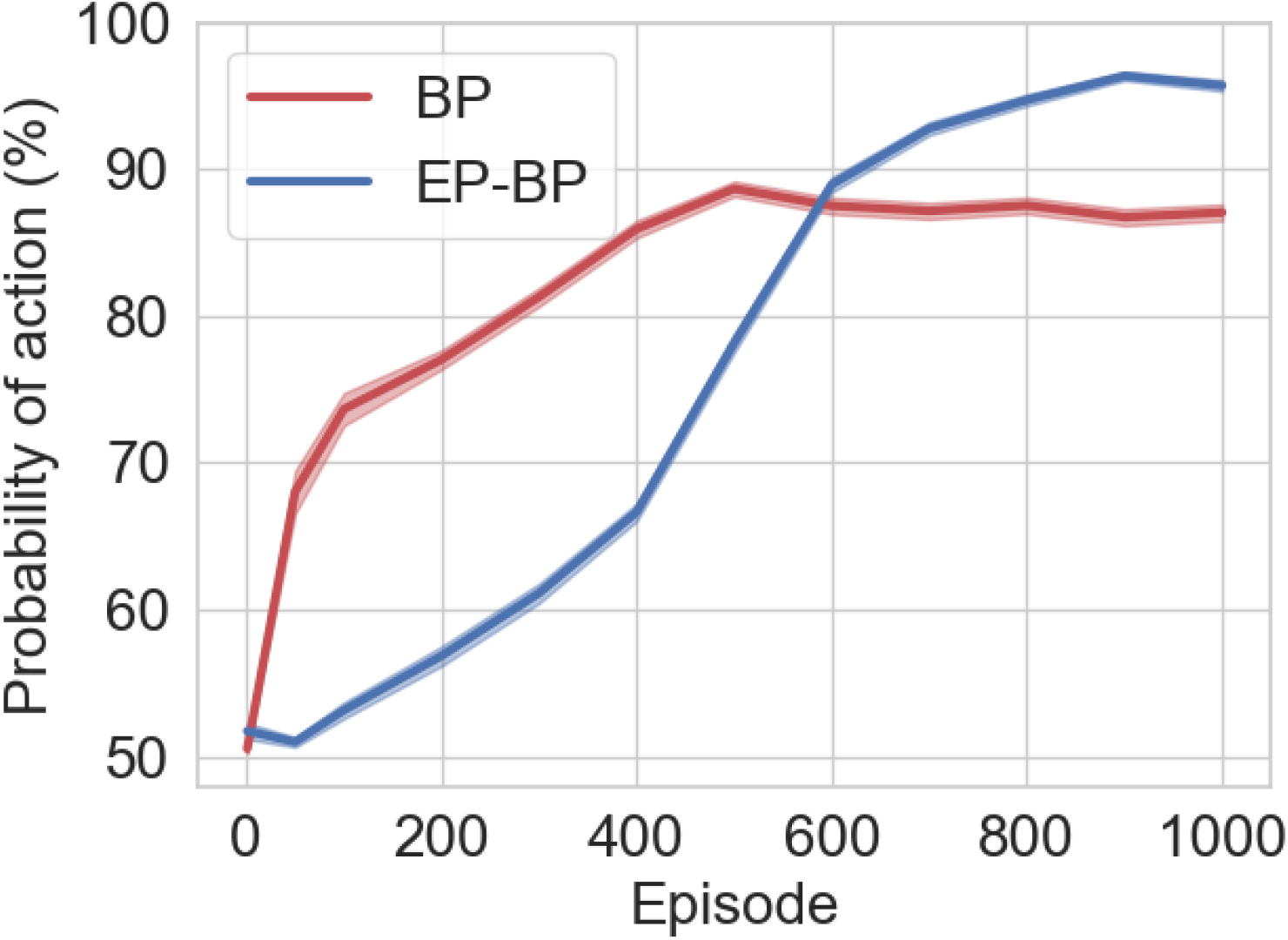
Mean and std error (SEM) (shaded area) for the probability of actions that EP-BP and BP models takes on CartPole-v0.

## Discussion

This study has explored the value of Equilibrium Propagation (EP) in reinforcement learning by proposing an Actor-Critic model with the Actor network trained by EP and the Critic network trained by backpropagation. The resulting models learn more consistent high-reward behavior than a baseline model trained exclusively by backpropagation. EP has been previously applied to image classification, but to our knowledge this is the first attempt to formulate an EP-based reinforcement learning system. Thus, we consider it to be an important development toward the next generation of biologically-plausible algorithms. Other, future developments should include application of EP to tasks like video classification [32] and speech recognition [33].

Further future work should also include a reinforcement learning system trained exclusively by EP, including a convolutional network for application to more complex tasks such as Atari games [34] or for neuronal data analysis tasks [35-38]. Another avenue for exploration would be the inclusion of neural adaptation [39, 40] ; a biologically-inspired modification to EP which previous work has shown to work well on image classification tasks, and may have value in reinforcement learning as well.

On three tasks investigated here, our EP-BP model works better than the AC trained only by BP. One of the reasons why it works better is, again, the higher probabilities of action. In the last phase of the training, we could see that our model is very stable and has higher probabilities of action. This means our model has enough information about the environment of the tasks, and the model takes optimal actions. This might happen because our model learns the environment slowly (Figures 1 and 5 tell us that our model learns the environment slowly), and this might lead the model to explore the environment to gather enough information and get the higher confidence of action.

## References

1. Rumelhart, D.E., G.E. Hinton, and R.J. Williams, Learning representations by back-propagating errors. nature, 1986. 323(6088): p. 533–536.

2. Silver, D., et al., Mastering the game of Go with deep neural networks and tree search. nature, 2016. 529(7587): p. 484–489.

3. Mnih, V., et al., Human-level control through deep reinforcement learning. nature, 2015. 518(7540): p. 529–533.

4. Goodfellow, I.J., J. Shlens, and C. Szegedy, Explaining and harnessing adversarial examples. arXiv preprint 1412.6572, 2014.

5. Williams, R.J., Simple statistical gradient-following algorithms for connectionist reinforcement learning. Machine learning, 1992. 8(3): p. 229–256.

6. Sutton, R.S. and A.G. Barto, Reinforcement learning: An introduction. 2018: MIT press.

7. Chung, S., An Alternative to Backpropagation in Deep Reinforcement Learning. 2020.

8. Pozzi, I., S. Bohte, and P. Roelfsema, Attention-Gated Brain Propagation: How the brain can implement reward-based error backpropagation. Advances in Neural Information Processing Systems, 2020. 33: p. 2516–2526.

9. Joel, D., Y. Niv, and E. Ruppin, Actor–critic models of the basal ganglia: New anatomical and computational perspectives. Neural networks, 2002. 15(4-6): p. 535–547.

10. Sheikhnezhad Fard, F., Modelling Human Target Reaching using A novel predictive deep reinforcement learning technique. 2018.

11. Takahashi, Y., G. Schoenbaum, and Y. Niv, Silencing the critics: understanding the effects of cocaine sensitization on dorsolateral and ventral striatum in the context of an actor/critic model. Frontiers in neuroscience, 2008. 2: p. 14.

12. Chalmers, E. and A. Luczak, Reinforcement Learning with Brain-Inspired Modulation can Improve Adaptation to Environmental Changes. arXiv preprint 2205.09729, 2022.

13. Ernoult, M., et al., Updates of equilibrium prop match gradients of backprop through time in an RNN with static input. Advances in neural information processing systems, 2019. 32.

14. Laborieux, A., et al., Scaling equilibrium propagation to deep convnets by drastically reducing its gradient estimator bias. Frontiers in neuroscience, 2021. 15: p. 129.

15. O’Connor, P., E. Gavves, and M. Welling. Training a spiking neural network with equilibrium propagation. in The 22nd international conference on artificial intelligence and statistics. 2019. PMLR.

16. Scellier, B. and Y. Bengio, Equilibrium propagation: Bridging the gap between energy-based models and backpropagation. Frontiers in computational neuroscience, 2017. 11: p. 24.

17. Scellier, B. and Y. Bengio, Equivalence of equilibrium propagation and recurrent backpropagation. Neural computation, 2019. 31(2): p. 312–329.

18. Almeida, L.B., A Learning Rule for Asynchronous Perceptrons with Feedback in a Combinatorial Environment, in Proceedings of the IEEE First International Conference on Neural Networks San Diego, CA, M. Caudil and C. Butler, Editors. 1987. p. 609–618.

19. Baldi, P. and F. Pineda, Contrastive learning and neural oscillations. Neural computation, 1991. 3(4): p. 526–545.

20. Pineda, F.J., Generalization of back-propagation to recurrent neural networks. Physical review letters, 1987. 59(19): p. 2229.

21. LeCun, Y., et al., Gradient-based learning applied to document recognition. Proceedings of the IEEE, 1998. 86(11): p. 2278–2324.

22. Krizhevsky, A. and G. Hinton, Learning multiple layers of features from tiny images. 2009.

23. Luczak, A., B.L. McNaughton, and Y. Kubo, Neurons learn by predicting future activity. Nature Machine Intelligence, 2022: p. 1–11.

24. Lin, L.-J., Reinforcement learning for robots using neural networks. 1992: Carnegie Mellon University.

25. Mnih, V., et al., Playing atari with deep reinforcement learning. arXiv preprint 1312.5602, 2013.

26. Wang, Z., et al., Sample efficient actor-critic with experience replay. arXiv preprint 1611.01224, 2016.

27. Wilson, M.A. and B.L. McNaughton, Reactivation of hippocampal ensemble memories during sleep. Science, 1994. 265(5172): p. 676–679.

28. Brockman, G., et al., Openai gym. arXiv preprint 1606.01540, 2016.

29. Kingma, D.P. and J. Ba, Adam: A method for stochastic optimization. arXiv preprint 1412.6980, 2014.

30. Römer, M., et al., Temperature Control for Automated Tape Laying with Infrared Heaters Based on Reinforcement Learning. Machines, 2022. 10(3): p. 164.

31. Maroti, A., Rbed: Reward based epsilon decay. arXiv preprint 1910.13701, 2019.

32. Karpathy, A., et al. Large-scale video classification with convolutional neural networks. in Proceedings of the IEEE conference on Computer Vision and Pattern Recognition. 2014.

33. Malik, M., et al., Automatic speech recognition: a survey. Multimedia Tools and Applications, 2021. 80(6): p. 9411–9457.

34. Bellemare, M.G., et al., The arcade learning environment: An evaluation platform for general agents. Journal of Artificial Intelligence Research, 2013. 47: p. 253–279.

35. Luczak, A., et al., Multivariate receptive field mapping in marmoset auditory cortex. Journal of neuroscience methods, 2004. 136(1): p. 77–85.

36. Luczak, A. and N.S. Narayanan, Spectral representation—analyzing single-unit activity in extracellularly recorded neuronal data without spike sorting. Journal of neuroscience methods, 2005. 144(1): p. 53–61.

37. Ponjavic-Conte, K.D., et al., Neural correlates of auditory distraction revealed in theta-band EEG. Neuroreport, 2012. 23(4): p. 240–245.

38. Ryait, H., et al., Data-driven analyses of motor impairments in animal models of neurological disorders. PLoS biology, 2019. 17(11): p. e3000516.

39. Kubo, Y., E. Chalmers, and A. Luczak, Biologically-inspired neuronal adaptation improves learning in neural networks. arXiv preprint 2204.14008, 2022.

40. Luczak, A. and Y. Kubo, Predictive neuronal adaptation as a basis for consciousness. Frontiers in Systems Neuroscience, 2021. 15.

